# paraCell: A novel software tool for the interactive analysis and visualization of standard and dual host-parasite single cell RNA-Seq data

**DOI:** 10.1101/2024.08.29.610375

**Authors:** Edward Agboraw, William Haese-Hill, Franziska Hentzschel, Emma Briggs, Dana Aghabi, Anna Heawood, Clare R. Harding, Brian Shiels, Kathryn Crouch, Domenico Somma, Thomas D. Otto

## Abstract

Advances in sequencing technology have led to a dramatic increase in the number of single-cell transcriptomic datasets available. In the field of parasitology these datasets typically describe the gene expression patterns of a given parasite species under specific experimental conditions, in specific hosts or tissues, or at different life-cycle stages. However, while this wealth of available data represents a significant resource for further research, the analysis of these datasets often requires significant computational skills, preventing a considerable proportion of the parasitology community from meaningfully incorporating existing single-cell data into their work. Here, we present paraCell, a novel software tool that automates the advanced analysis of published single-cell data without requiring any programming ability. On our free web-server, we demonstrated how to visualise data, re-analyse published *Plasmodium* and *Trypanosoma* datasets, and present novel *Toxoplasma*-mouse and *Theileira*-cow atlases to study the impact of IFN-γ and host genetic susceptibility.

## Background

Single-cell transcriptomics sequencing (scRNA-seq) was first introduced in 2009(Tang, et al., 2009) and has since had a dramatic impact on the investigation of cellular heterogeneity, the identification of novel cell types, and the discovery of new drug targets (Jovic, et al., 2022). It has also enabled projects such as the Human Cell Atlas, which seeks to catalogue and describe every cell type in the human body(Rozenblatt-Rosen, et al., 2017).

Single-cell sequencing has enabled similar “cell atlas” projects in the field of parasitology, typically detailing the full life cycle of various parasite species such as *Plasmodium* (Howick, et al., 2019), *Toxoplasma* (Xue, et al., 2020) or *Schistosoma* (Diaz Soria, et al., 2020). Other applications include the in-depth description of specific life-stages (Briggs, et al., 2021) or sexual development (Hentzschel, et al., 2022) in parasites, in addition to the investigation of host-parasite interactions which can reveal differential host immune responses and parasite invasion preferences (Greenwood, et al., 2016).

Host-parasite interaction analysis is enabled by dual RNA-seq, a novel transcriptomic approach capable of capturing the gene expression profiles of both a given cell and any infectious agents within (Greenwood, et al., 2016). This technique is particularly useful in the study of intracellular parasites such as *Theileria*, which is known to induce an oncogenic state in infected bovine leukocytes (Jensen, et al., 2009), *Toxoplasma* which secretes hundreds of proteins into host cells, altering the host transcriptome and is responsible for toxoplasmosis (Rastogi, et al., 2020) or *Plasmodium*, the causative agent behind malaria (Hentzschel, et al., 2022) . In cases such as this, dual scRNA-seq can be used to capture both host and parasite mRNA transcripts from infected cells, enabling the investigation of host-parasite interactions on the single-cell level and allowing the complementary interrogation of critical processes such as infection, pathogenesis and cellular immunity. However, there is a lack of dedicated tools for the analysis of host-parasite dual scRNA-seq datasets (Karagiannis, et al., 2023).

Single-cell datasets are inherently collaborative projects, relying on both effective cell culturing or sample collection by biologists in the wet lab and valid statistical analysis by bioinformaticians in silico. This statistical analysis of single-cell transcriptomic data is typically handled by general purpose packages such as Seurat (Butler, et al., 2018) or Scanpy (Wolf, et al., 2018), often in combination with more specific tools and/or software packages designed for specific steps in the single-cell analysis pipeline, such as cluster annotation, differential gene expression analysis or trajectory inference (Slovin, et al., 2021).

However, this computational analysis is most often the sole responsibility of a trained bioinformatician, with the other, wet lab-based members of the collaboration having relatively restricted access to the process. This can be a limiting factor in bioinformatic research, as the in-depth biological knowledge that wet lab-based collaborators possess could provide the key insights needed to guide the analysis.

Cell atlases (such as the Human Cell Atlas, Malaria Cell Atlas (Howick, et al., 2019), cell atlases made available via VEuPathDB (Alvarez-Jarreta, et al., 2024) and SchistoCyte Atlas (Wendt, et al., 2021)) help the field of single-cell transcriptomics overcome this analysis bottleneck, facilitating more effective collaboration between biologists and bioinformaticians by visualizing single-cell datasets via single-cell browsers. Single-cell browsers are software tools that automate the visualization of single-cell data, providing users with interactive graphical interfaces that remove the requirement for computational skills ordinarily needed to interact with scRNA-seq data. These platforms help solve a common problem in the multi-disciplinary teams characteristic of bioinformatics research - a disconnect between the experimental scientists who generate data and the computational scientists who analyse it (Morrison-Smith, et al., 2022) - by providing a single, shared representation of a given dataset around which to base discussion.

Popular single-cell browsers include the UCSC Cell Browser (Speir, et al., 2021), the Single Cell Explorer (Feng, et al., 2019), and CELLxGENE (Megill, et al., 2021), a Chan Zuckerberg Initiative project that was recently identified as the premiere option for the publication of single-cell data (Cakir, et al., 2020) and that is used to visualize data sets hosted on the Human Cell Atlas. CELLxGENE is distinguished not only by its fully comprehensive user interface and efficient implementation (which allows for the visualization of millions of cells (Megill, et al., 2021)), but also by its inclusion of in-built options such as differential gene expression (DGE) analysis and marker gene identification; a notable departure from other single-cell browsers which typically focus exclusively on visualization (Table 1). CELLxGENE is used in VEuPathDB (Alvarez-Jarreta, et al., 2024) for the representation of single cell data from parasites and their hosts… However, the focus of CELLxGENE still leans more towards visualization than analysis.

**Table 1:** Comparison of paraCell to other available single-cell browsers (Cakir, et al., 2020).

|  |  |  |  |  |  |  |
| --- | --- | --- | --- | --- | --- | --- |
|  | UCSC<br>Cell<br>Browser | Single<br>Cell<br>Explorer | iSEE | CELLx<br>GENE | CELLx<br>GENE<br>VIP | paraCell |
| Web Sharing | ✓ | ✓ | ✓ | ✓ | ✓ | ✓ |
| Interactivity | ✓ | ✓ | ✓ | ✓ | ✓ | ✓ |
| Cell Select Functionality | ✓ | ✓ | ✓ | ✓ | ✓ | ✓ |
| Multiple Embeddings |  | ✓ | ✓ | ✓ | ✓ | ✓ |
| DEG Analysis |  |  | ✓ | ✓ | ✓ | ✓ |
| Additional Visualization Options |  |  |  |  | ✓ | ✓ |
| Additional Search Options |  |  |  |  |  | ✓ |
| External Database Links |  |  |  |  |  | ✓ |
| Trajectory Inference |  |  |  |  |  | ✓ |
| Host-Parasite Interaction Analysis |  |  |  |  |  | ✓ |
| Docker |  |  |  | ✓ |  | ✓ |

The third-party plugin CELLxGENE VIP seeks to overcome this limitation by providing users with additional options for the analysis and visualization of single-cell data (Li, et al., 2022), such as heatmaps, violin plots and volcano plots.

However, while VIP significantly expands the number of analysis options available within the CELLxGENE framework, it is still limited. VIP does not provide users with the ability to search a given dataset using gene product names or GO terms and does not include any provision for trajectory inference, enrichment analysis after differential expression, nor the interrogation of the dual scRNA-seq datasets required for host-parasite interaction analysis; functionalities which would be highly appreciated in the field of parasitology. In this way CELLxGENE VIP is very similar to the other actively maintained cell atlas services, none of which provide all these options (Table 1).

Here we present paraCell, a novel software tool based on CELLxGENE and VIP, designed for the visualization and analysis of single-cell parasitological data. Example paraCell atlases are available at http://cellatlas.mvls.gla.ac.uk. Five use cases are detailed in this paper, including two novel dual scRNA-seq host-parasite datasets describing the impact of Theileria infection on either *Bos indicine* or *Bos taurine* cattle, and *Toxoplasma*-infected bone marrow-derived macrophages treated with IFN-γ.

## Results

We implemented paraCell as an expansion of CELLxGENE and VIP by adding new analysis, visualization, and search options to the application. We also increased interoperability between the plugin and base CELLxGENE as well as external database systems such as NCBI (Clough, et al., 2024) and VEuPathDB (Figure 1).

**Figure 1:**
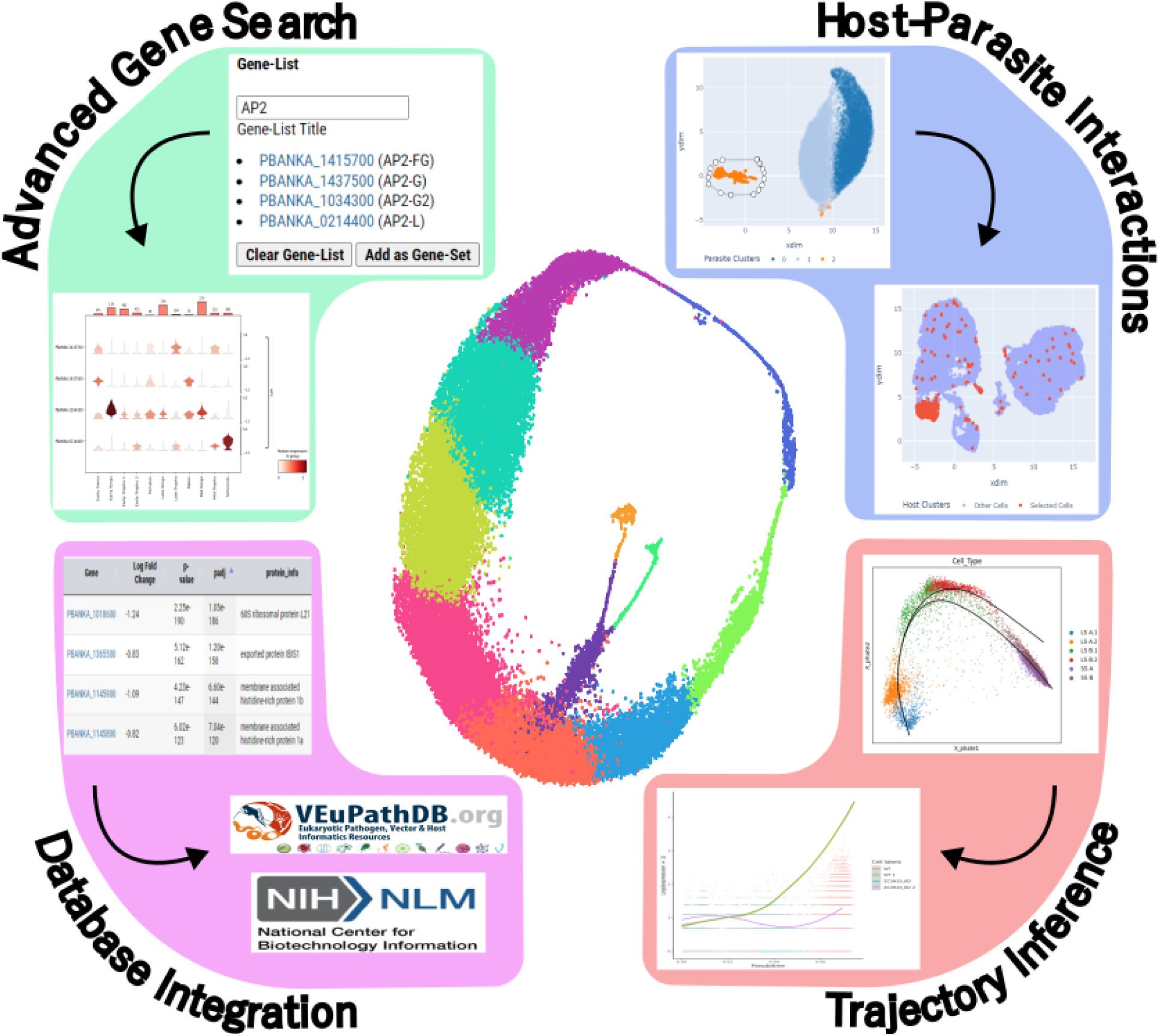
Overview of the four main options in paraCell. Advanced Gene Search - a custom gene-set is used to create a VIP multi-gene violin plot (Plasmodium berghei dataset (Hentzschel, et al., 2022)). Host-Parasite Interactions – a Selection made on the Parasite UMAP is used to update the Host UMAP (Cow-Theileria dataset). Database Integration - a paraCell results table incorporates links to relevant external database systems (P. berghei dataset (Hentzschel, et al., 2022)). Trajectory Inference - Slingshot trajectories are drawn over PHATE space of single-cell data, enabling tradeSeq to plot the expression of a gene of interest across each lineage, split by the levels of a specified experimental condition (Trypanosoma brucei dataset (Briggs, et al., 2021)).

The installation of paraCell’s dependencies is enabled by the “paraCell” Conda environment provided alongside the main application. Once installed, paraCell facilitates the transformation of AnnData objects (Wolf, et al., 2018) into either local or online cell atlases.

Like CELLxGENE and VIP, paraCell is based on the AnnData object – a standard and highly flexible file format capable of holding all the diverse information associated with a given scRNA-seq experiment, from gene expression counts to cell annotations. This adaptability lets the AnnData input file provided to paraCell incorporate “paraCell_setup”, a Python dictionary which controls the activation of paraCell’s new functionality. Further information on the creation and use of “paraCell_setup” can be found in the “Wiki” section of the paraCell GitHub repo. We have an instance running on http://cellatlas.mvls.gla.ac.uk/ that visualises several datasets and which was used to generate the five use-cases below.

### Comparison to other available tools

CELLxGENE VIP extends the CELLxGENE framework, and paraCell builds on CELLxGENE VIP, resulting in a platform distinguished by an intuitive user interface and a comprehensive suite of additional analysis and visualization options (Table 1). paraCell benefits from active development in the CELLxGENE community and is further differentiated from the alternatives by its focus on parasitology, providing functionality such as links to the popular parasite data warehouse VEuPathDB, additional annotation options in differential expressed gene (DEG) analysis and provisions for host-parasite interaction analysis.

The following four sections demonstrate the unique capabilities of paraCell using case studies based on parasite scRNA-seq datasets (Figure 1). The specific paraCell workflow associated with each case study can be found in the paraCell GitHub page.

### Case Study 1 Advanced Search Options

A common first step taken by parasitologists when interacting with a single-cell dataset is to check whether any cells express a specific gene of interest. Although base CELLxGENE does include a search bar, this native search functionality is limited to a single identifier, typically the gene ID or gene name. paraCell seeks to overcome this limitation by implementing Advanced Search Options (Figure 1, top left).

paraCell implements an additional search bar, capable of utilizing both colloquial gene names and gene product descriptions as identifiers. In-built autocomplete functionality also enables the use of gene “terms’’ as identifiers. For example, in Figure 1 we used the term AP2, which is an important transcription factor family in *Plasmodium*, and other important apicomplexans, to identify all of the AP2 related genes in the dataset. Users can then save specific items retrieved via this search bar to a custom gene set, enabling the generation of a variety of informative plots via CELLxGENE VIP (e.g. violin plots depicting the expression of each gene within specific groupings of cells).

paraCell also implements a Gene Ontology (GO) search bar, capable of retrieving all the genes associated with a specific GO term and saving the retrieved items as a gene set for later use with CELLxGENE VIP analysis and/or plotting functions.

### Case Study 2: DEG and Integration of External Databases

CELLxGENE includes native DEG analysis functionality, which lets users select and then compare two specific cell populations on the mRNA level. The top genes showing elevated expression in each population are saved and presented to the user as gene sets.

paraCell enables users to import these gene sets directly into the plugin. Any gene set available in paraCell can then be used to form a search strategy on a preset external database: currently available options are VEuPathDB or the NCBI gene database. The database associated with a given cell atlas is extracted from a list stored in the underlying annData object.

This approach empowers users to combine the functionality of CELLxGENE and paraCell with that of external databases, such as the Gene Ontology Analysis option built into VEuPathDB.

The utility of paraCell’s interconnectivity is demonstrated here by the re-analysis of scRNA-seq *Plasmodium berghei* data produced by Hentzschel et al (Hentzschel, et al., 2022). The original study charted the sexual development of *Plasmodium* parasites within host cells of varying maturity. Following this, a paraCell atlas linked to the PlasmoDB database (Aurrecoechea, et al., 2009) was created to present the dataset to the public (https://cellatlas-cxg.mvls.gla.ac.uk/Plasmodium_berghei-Multi.Tissue/) and used to extract novel insights regarding the early stages of *Plasmodium* gametogenesis (Figure 2).

**Figure 2:**
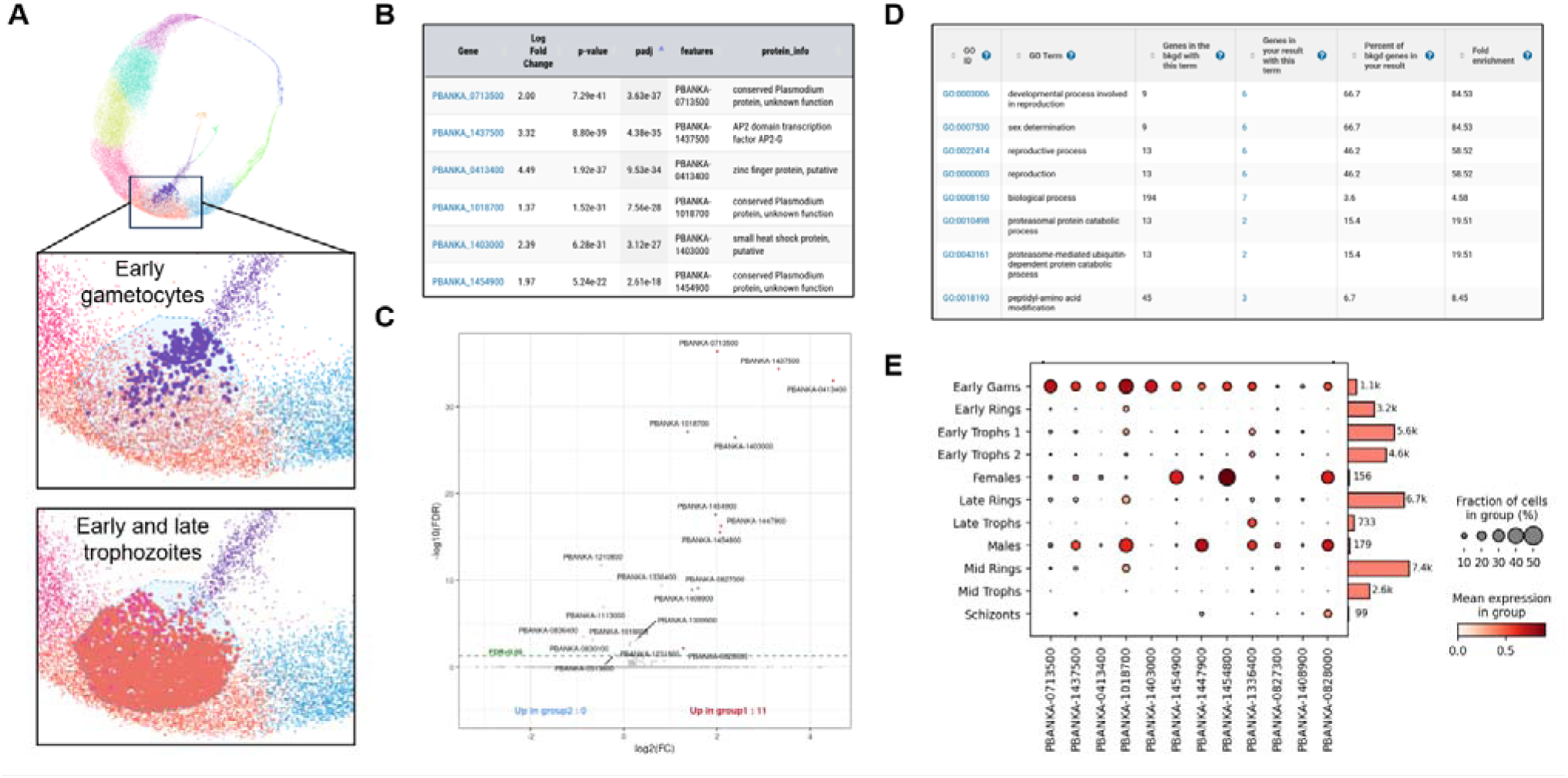
A) The selection of two cell populations (early gametocytes vs. early/late trophozoites) in CELLxGENE. B) DEG analysis results table (early gametocytes vs. early/late trophozoites). C) Volcano plot of the DE genes. D) Gene Ontology enrichment results table for the DE gene-set. E) Dot Plot showing the relative expression level of each DE Gene in each stage of the malaria life cycle.

Base CELLxGENE functionality was used to target the intersection between the trophozoite stage and the early gametocytes (Figure 2A). CELLxGENE VIP functionality was then used to perform differential gene expression analysis between trophozoite and early gametocyte cells within this intersection (Figure 2B).

Several genes known to characterize *Plasmodium* gametogenesis, such as AP2G (PBANKA_1437550), gametocyte development gene 1 (GDV1, PBANKA_0828000) and male development protein 2 (MD2, PBANKA_1447900) and 3 (MD3, PBANKA_0413400), were found to be upregulated in gametocytes. In line with this, exporting the set of DE genes to PlasmoDB via paraCell for GO analysis revealed an enrichment of terms related to sexual determination as well as reproductive and developmental processes (Figure 2D).

Notably, this analysis also identified several novel genes upregulated in gametocytes but not previously associated with *Plasmodium* gametogenesis, such as PBANKA_0713500, a conserved protein of unknown function which was the most significant hit of all DEG, or PBANKA_1403000, a putative small heat shock protein. These genes were found to be primarily expressed in gametocytes, both early in development as well as in mature male and female cells (Figure 2E).

These results demonstrate the synergy paraCell promotes between CELLxGENE, CELLxGENE VIP, and external databases (e.g. PlasmoDB). Utilizing the functionality of all three systems enables a highly targeted, interactive analysis which has identified new candidate genes potentially implicated in the early stages of *Plasmodium* gametogenesis, all without any need for programming experience on the part of the user.

### Case Study 3: Trajectory Inference and Trajectory-based Differential Expression Analysis in paraCell

paraCell expands the functionality of VIP to include both trajectory inference and trajectory based differential expression analysis. The *Trajectory Inference* tab can display precomputed Slingshot (Street, et al., 2018) trajectories, as well as generate partition-based graph abstraction (PAGA) (Wolf, et al., 2019) maps describing the overall topology and connectivity of annotated scRNA-seq data. And the *tradeSeq* (Van den Berge, et al., 2020) tab implements the eponymous tool, an R analysis package that enables trajectory based differential expression analysis. This tab is capable of both displaying pre-computed tradeSeq results and generating new tradeSeq plots indicating the relationship between the expression of a given gene and the progression of a specified lineage.

Together these features allow paraCell atlases to extract and display a wider range of information from accompanying papers and empower users to explore scRNA-seq datasets in new ways.

The utility of these features is demonstrated here by the re-analysis of scRNA-seq data generated and published by Briggs et al (Briggs, et al., 2021). The original study described the development of bloodstream form *Trypanosoma brucei* parasites from a replicative ‘slender’ form to a transmissible ‘stumpy’ form in both a wildtype (WT) and a ZC3H20 knockout (KO) population, where the deletion of this critical RNA-binding protein prevents stumpy formation (Briggs, et al., 2021). Two trajectories were identified via Slingshot - a complete trajectory describing the successful differentiation of WT parasites, and a truncated trajectory describing the incomplete differentiation of KO parasites (Figure 3). Genes significantly associated with each trajectory were then identified via tradeSeq.

**Figure 3:**
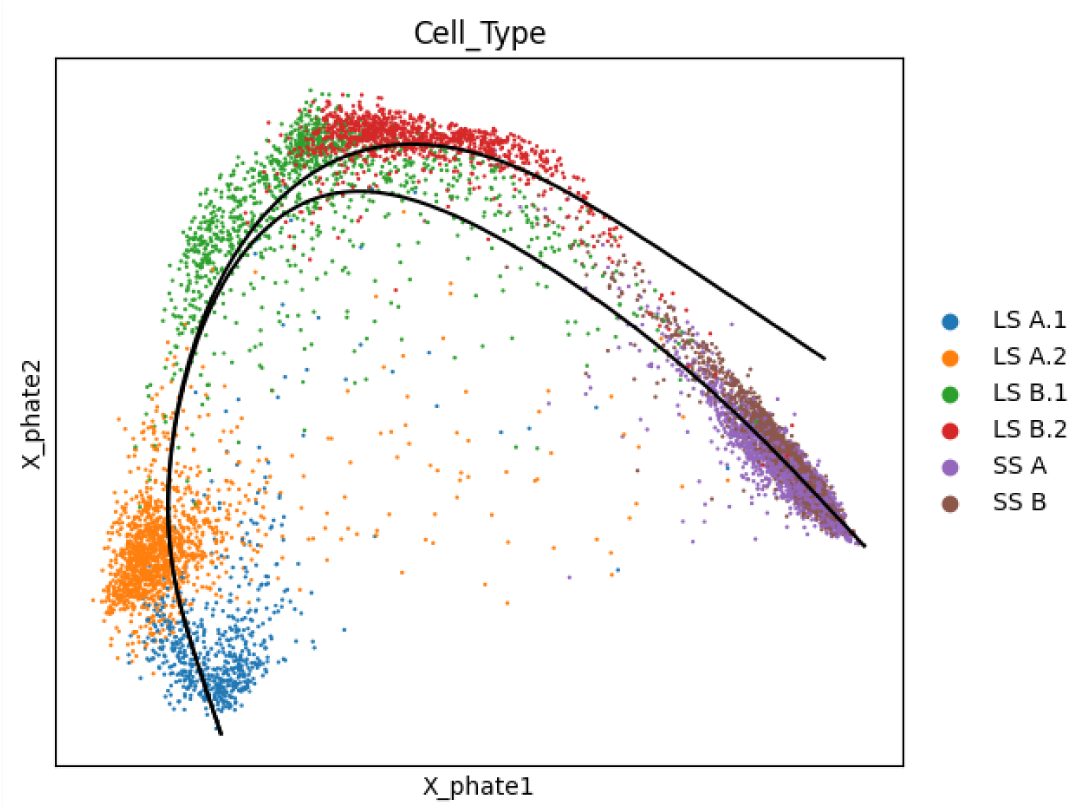
Low dimensional plot (PHATE) of the T. brucei scRNA-seq dataset, grouped by cell type, as presented by the Trajectory Inference Tab in paraCell. The branched trajectories identified in the data are indicated by black lines.

Following this a paraCell atlas was created to present the results to the public (https://cellatlas-cxg.mvls.gla.ac.uk/Trypanosoma_brucei-slender2stumpy/), and the *tradeSeq* tab was used to identify new genes associated with the successful differentiation of *T. brucei* parasites.

The search for genes differentially expressed across only the WT trajectory from slender to stumpy forms, and not differentially expressed during the KO trajectory, implicated 385 genes encoding hypothetical proteins linked to slender to stumpy life cycle development but currently uncharacterised. Expression patterns of these genes were then plotted using paraCell, allowing further investigation. Here we discuss one gene of interest revealed by this approach, Tb927.6.630. The tradeSeq plot generated for Tb927.6.630 revealed that the expression of the gene peaks near the endpoint of the WT trajectory, a region corresponding to the transmissible stumpy form of the parasite (Figure 4A). Mining other available *T. brucei* resources identified that the protein encoded by Tb927.6.630 localised to the nucleus of bloodstream and procyclic form parasites (Moloney, et al., 2023), and specifically the nucleolus of procyclic form parasites (Figure 4B), as revealed by fluorescent protein tagging project TrypTag (Billington, et al., 2023). This protein is predicted to have a complex structure (Figure 4C) that, when aligned to the database of all available protein structures via FoldSeek (van Kempen, et al., 2023), returns uncharacterized homolog proteins in related trypanosomatids such as *Trypanosoma cruzi* (Tc00.1047053510535.10) and *Leishmania infantum* (LINF_220022400), as well as a partial alignment to the *Arabidopsis thaliana* protein Katanin p80 WD40 repeat-containing subunit B1 homolog (KTN80.1), which has predicted functions in rapid reorganisation of cellular microtubules (Wang, et al., 2017) (Figure 4D). KTN80.1 is the non-catalytic subunit of the microtubule-severing enzyme complex katanin (Wang, et al., 2017).

**Figure 4:**
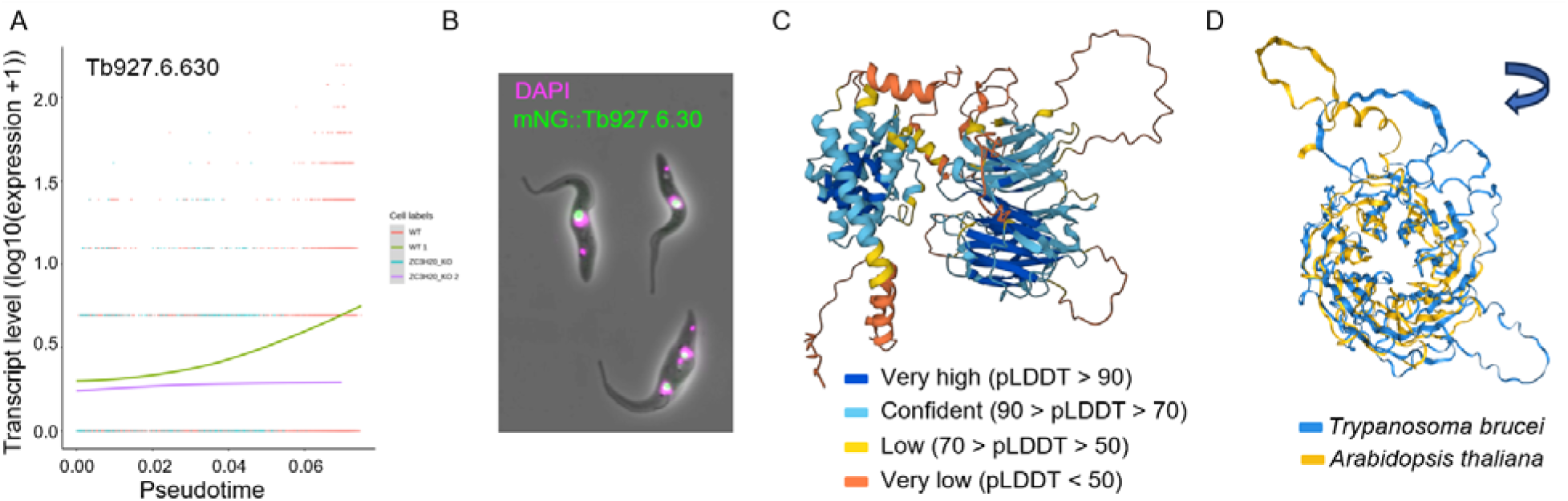
Investigation of a novel life cycle regulated Trypanosoma brucei protein. A) mRNA level (y-axis) changes for Tb927.6.630 across pseudotime (x-axis). mRNA count for individual wild type (WT, red) and ZC3H20 null mutant (ZC3H20 KO, teal) Trypanosoma brucei, during life cycle progression from slender to stumpy forms. Trajectory 1 of wild type parasites (WT 1, green) and alternative trajectory 2 taken by ZC3H20 mutants (ZC3H20 KO 2, purple) show differential expression patterns during full development to stumpy forms in wild type parasites and truncated development of mutant parasites. B) Localisation of NeonGreen endogenously N-terminal tagged Tb927.6.630 protein (mNG: Tb927.6.630, green) to the nucleolus of procyclic form parasites. Nuclei visualised by DAPI staining DNA (pink). Image retrieved from TrypTag database resource (Billington, et al., 2023). C) Predicted protein structure, coloured by per-residue confidence score (pLDDT). Image retrieved from the AlphaFold Protein Structure Database. D) Protein alignment of the WD40 repeat-containing domain of T. brucei Tb927.6.630 and Arabidopsis thaliana protein KTN80.1.

Tb927.6.630 is predicted to contain seven WD40 repeat regions which together form a circularized β-propeller. A similar structure has been documented in KTN80.1 and across the homologous p80 proteins in animal cells where the WD40 domain targets katanin to the centrosome (Roll-Mecak and McNally, 2010). Together this information highlights a complex protein that may have yet unexplored functions relating the *T. brucei* life cycle development, likely via structural changes to nuclear microtubules.

This analysis was only possible with paraCell given the ability to visualise and analyse the trajectory (trajectory inference and tradeSeq tabs). In summary, we findings highlight the utility of the, demonstrate how the additional options offered by paraCell facilitate the re-analysis of published data and, by doing so, encourage the generation of new biological insights.

### Case Study 4: Annotating and Analyzing *Host-parasite data 1*

The Host-Parasite Interactions (HPI) tab enables the analysis and visualization of dual scRNA-seq host-pathogen datasets. These datasets are important for exploring intracellular pathogens and are characterized by each cell containing both host and parasite genes. paraCell allows users to subset and visualize the underlying data object, facilitating the comparison of host and parasite behaviour at the gene expression level. This functionality is demonstrated first with a *Toxoplasma*-Mouse atlas, and secondly with a *Theileria*-Cow atlas.

To study *Toxoplasma*-Mouse interactions we extracted and differentiated murine bone marrow derived macrophages (BMDMs) which were unstimulated or stimulated with IFN-γ and infected with type 1, RH *Toxoplasma gondii*.

After 10x Chromium with Illumina sequencing, we obtained 1600, 1400 cells, with 3900 and 3100 mouse genes detected per cell and around 950 and 850 parasite gene, for the WT and IFN-γ stimulated infected cells, respectively. Data were processed using Seurat in R (see methods) and after filtering we had 1407 and 1258 cells for the treated and untreated. The R object was converted to an “AnnObejct” and loaded into paraCell (https://cellatlas-cxg.mvls.gla.ac.uk/Toxoplasma_gondii-murine.bone.marrow.derived.macrophages/).

Often, users receive well annotated objects to work with. However, here we show that paraCell can be used to identify clusters and to explore how parasites modulate host cell processes. The first analysis was to check the parasite frequency, showing that most clusters are host cells and we retrieved just around 100 parasite cells per run (Figure 5A). This was largely due to a low initial multiplicity of infection, and the comparatively low levels of parasite transcripts, especially when compared to highly active INF-activated macrophages.

**Figure 5:**
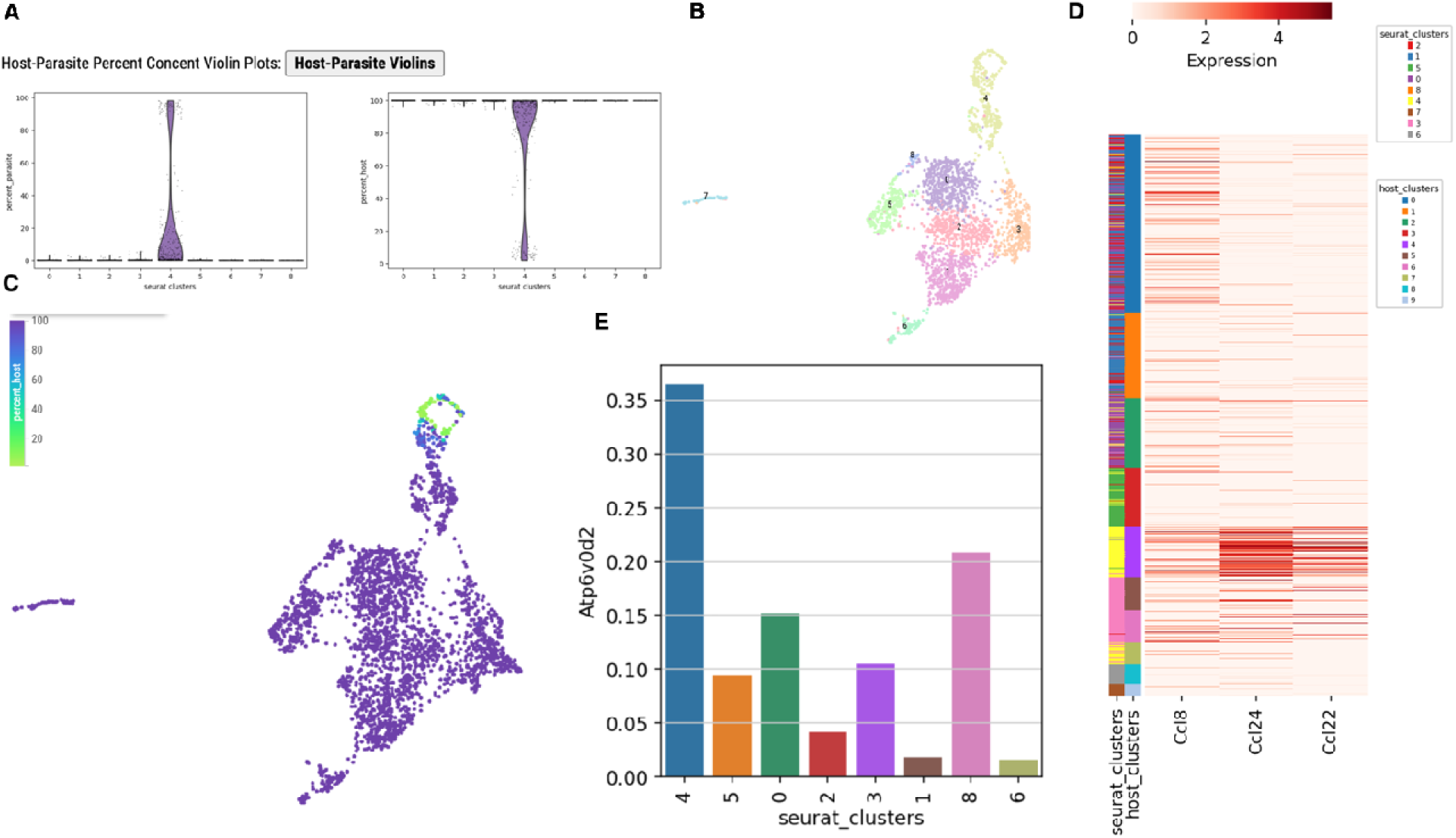
Overview of Toxoplasma-mouse cell atlas. A) Violin plot of percentage of parasite cells (left) and host cell (right) in the dataset. B) Clustered data in paraCell C) Showing the host content in the UMAP with one cluster being mostly parasite D) Expression of Atp6v0d2 E) Expression of key marker genes versus the Seurat and host clusters.

The effects of IFN stimulation could clearly be seen in the DEG analysis between runs, with an upregulation of IFN-γ-stimulated (e.g. Irf1, Gbp2, Gbp4) genes. From the combined UMAP, multiple macrophage populations could be identified (Figure 5B). Cluster 3 cells expressed a strong signature of IFN-γ-stimulation (high levels of Cxcl9, Cxcl10 and vim). Cluster 6 expressed markers for inflammatory monocytes (Ly6c2 and Ccr2), cluster 7 expressed markers of M2 macrophages (including Arg1, Chil3 and Retnla) and cluster 8 probably contained cycling macrophages, expressing multiple genes involved in DNA replication such as Smc4, Top2a, Pclaf and Rrm1.

Cluster 4 contains *T. gondii* infected macrophages (Figure 5C). These cells have upregulated Ccl22 and Ccl24(Figure 5D), as previously described(Gossner and Hassan, 2020). A novel marker genes was *Atp6v0d2*, which is highest expressed in that cluster (Figure 5E). Interestingly, this cluster also expresses Ccl8. Ccl8 (also called MCP-2) is a chemoattractant factor expressed on a range of inflammatory cells, but has not previously been associated with *T. gondii* infection. Ccl8 is upregulated in Mycobacteria infection(Liu, et al., 2013) and has recently been shown to be upregulated in response to lactate(Zhou, et al., 2023). *T. gondii* is known to secrete lactate into the host cell through FNT1(Zeng, et al., 2021), suggesting a mechanism for this host response.

The tools available allowed this analysis to be performed by a first-time user of single cell data with minimal training, highlighting the usability of the paraCell interface, and its ability to allow biological insights by the end users of the data.

### Case Study 5: Visualizing Host-Parasite Interactions in paraCell 2

As second host-parasite example we infected bovine macrophage cell lines from two cow strains with *Theileria annulata*. One cow breed (Holstein) is known to be susceptible to the parasite, while Sahiwal is resistant. We performed scRNA-Seq on the cells and obtained around 9000 cells, with ∼4000 genes per host cell and 480 vs 667 genes for the parasites (Holstein vs Sahiwal). Data were processed in R using Seurat (as above), leaving us with 8433 and 7831 cells for Holstein and Sahiwal, respectively. In contrast to the *Toxoplasma* example, using paraCell, we could see that most of the host cells were infected with parasites (Supplemental Figure 1, assessed via “host-parasite” violin plots option). Additionally, host and parasite marker genes can be identified for categorical annotation levels, showing a high percentage of parasites in clusters 4 and 6. These features provide users with an overview of host and/or parasite-specific gene expression within a dataset.

paraCell can identify different cell clusters (Figure 6A); based on marker genes (Figure 6B) some clusters are merged because they exhibit a similar macrophage state signature: clusters 0, 3, 8, and 11 are cycling macrophages expressing the marker MKI67 (Figure C); clusters 9 and 10 are M2-like macrophages expressing MMP9/COX2; clusters 2, 4, 5, and 12 are macrophages actively responding to the infection, expressing LGALS3 and TRAF3IP3; and clusters 6 and 7 are non-activated macrophages. The percent expression of host and/or parasite genes within categorical annotations can be assessed via “host-parasite” violin plots (Supplemental Figure 1). paraCell’s features enable users also to explore the relationship between host and parasite gene expression in an interactive and intuitive manner. In this case study, cells from Sahiwal cattle (*Bos indicus*), which are tolerant to the infection, were compared to cells from Holstein cattle (*Bos taurus*), which are susceptible to acute disease. Despite both cell types being infected by *Theileria annulata* and reaching a similar cycling stage, the integration analysis reveals that Sahiwal cells differentiate into an active macrophage cluster absent in Holstein cells (Figure 6B-C).

**Figure 6:**
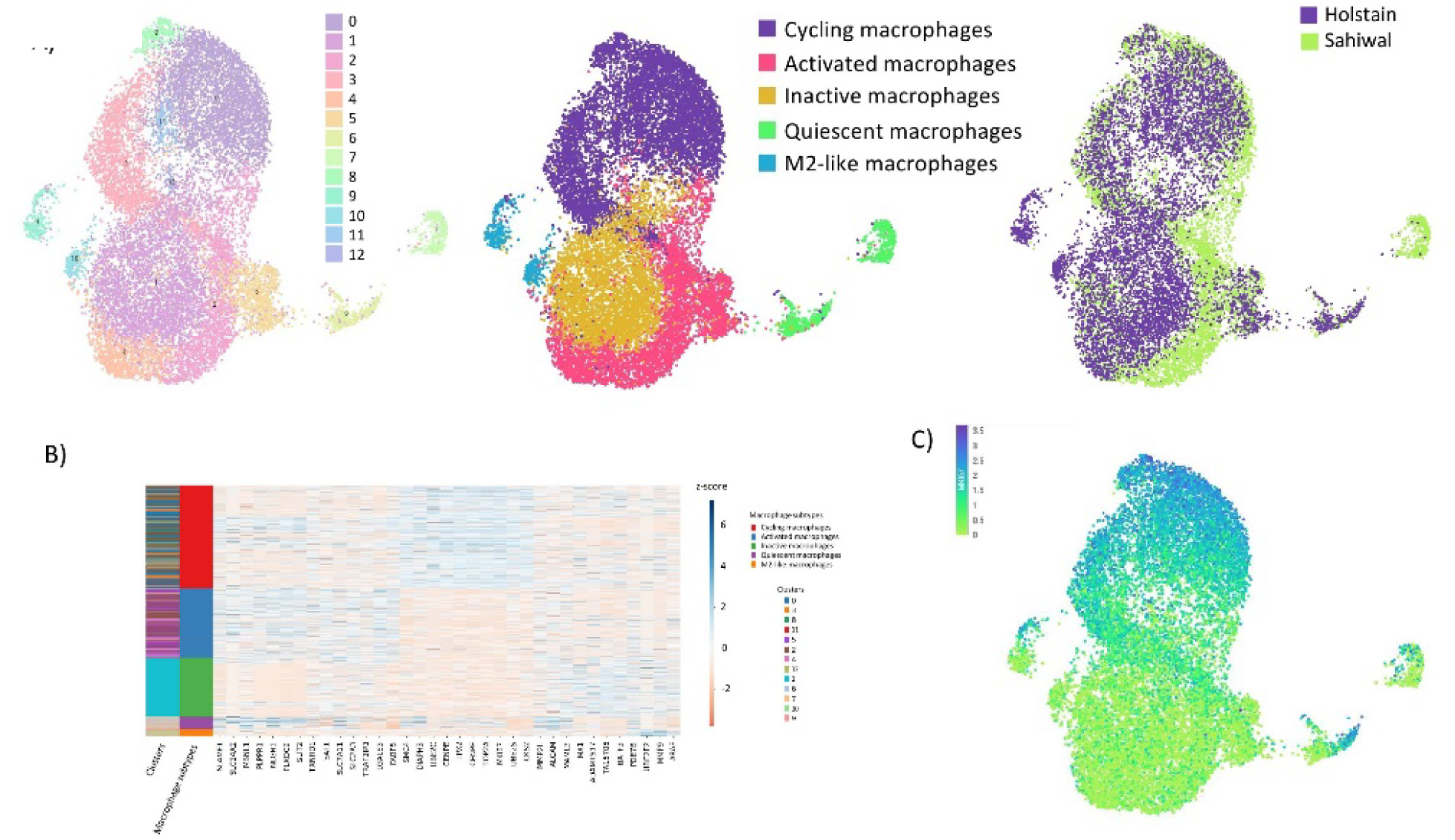
A) Cow-Theileria Combined UMAP, produced by reducing dimensions using both host and parasite genes in Seurat, showing clusters (left), macrophage subtypes (middle), and cattle species (right). B) Heatmap showing genes DE for each macrophage subtypes. C) MKI67 gene expression, as one of the key markers to identify macrophage in cell cycle stage.

We first selected and searched for differentially expressed genes between the cycling cells of the two cell lines (Figure 7A). The Gene Set Enrichment Analysis (GSEA) indicates an active interferon (IFN) response in the Sahiwal cells during the cell cycling stage, compared to Holstein cells (Figures 7B-C). We used the download option of paraCell and the results can be found in Supplementary Table 1). Conversely, the Holstein cells exhibit a gene signature indicative of a pro-carcinogenic profile, corroborating previous literature(Larcombe, et al., 2022).

**Figure 7:**
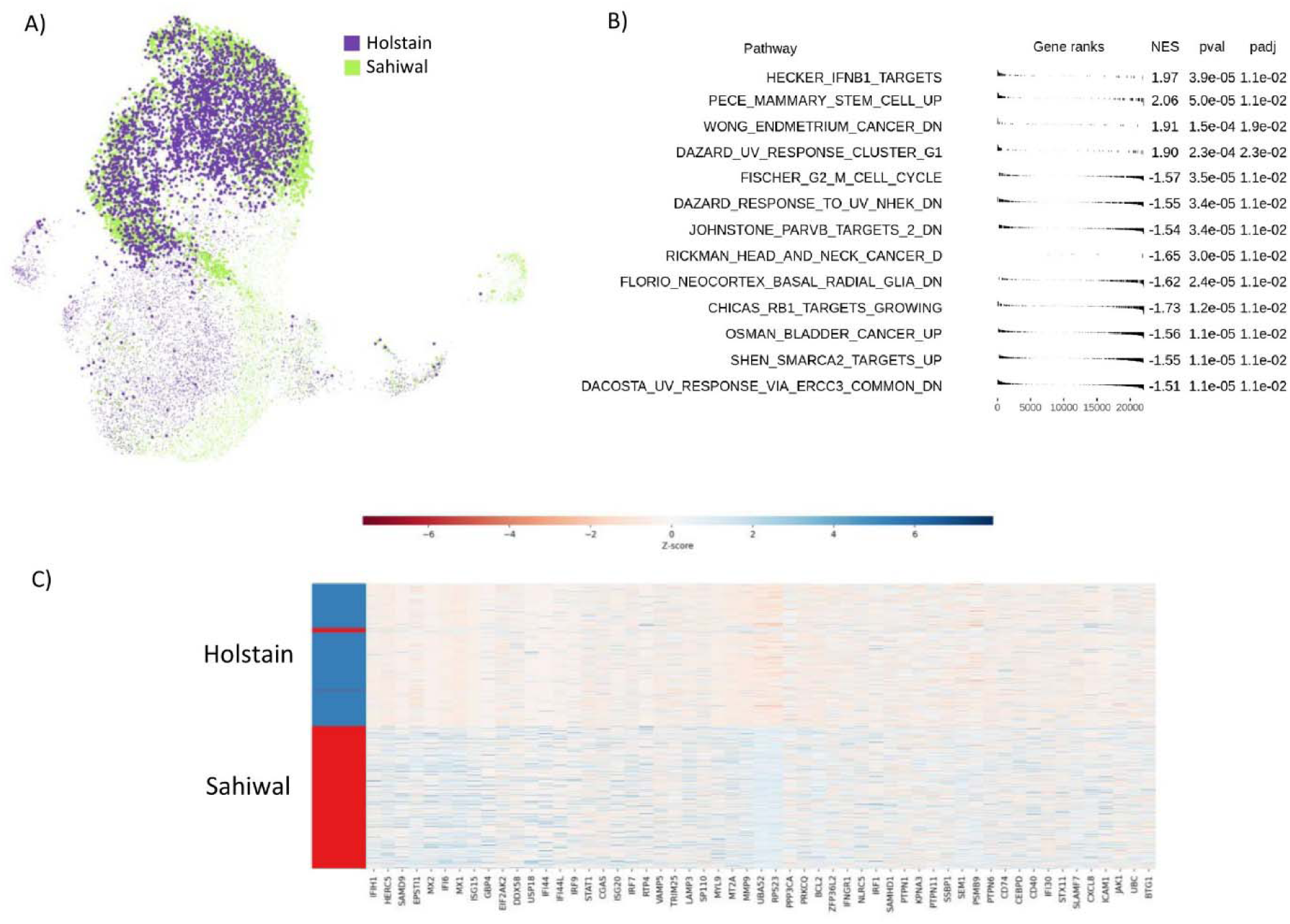
A) Cow-Theileria Combined UMAP, showing Holstain and Sahiwal clusters in violet and green respectively. B) GSEA showing pathway with >0 NES score, upregulated in Sahiwal, and <0 upregulated in Holstain C) Heatmap showing all the genes DE involved in at least one interferon pathway statistically significant for GSEA

The Host-Parasite UMAPs produced by paraCell are interactive Plotly graphs. The lasso select function in Plotly allows for freeform selection of cell subsets, dynamically updating graphs. Lasso selections made on the “source” graph (e.g., the Parasite UMAP) automatically update the “target” graph (e.g., the Host UMAP). Selected cell IDs on the source graph are extracted to either highlight the corresponding cells on the target graph or to produce a new version of the target graph displaying only the selected cells, as illustrated in Supplemental Figure 2.

In summary, paraCell provides a robust and interactive platform for the detailed exploration of host-parasite interactions at the single-cell level, offering valuable insights into the gene expression dynamics and cellular responses in infected populations.

## Discussion

In recent years, there has been a push for the adoption of findability, accessibility, interoperability, and reusability (FAIR) data-sharing principles in science (Barker, et al., 2022), which emphasize the utility of automated data retrieval and the importance of making published data easily available for re-use by the wider academic community.

However, the adoption of these principles has been slow in bioinformatics, hindered by the rapid pace of technological development and the frequent need for specialist knowledge during data analysis. Software tools and programming languages are frequently updated, resulting in compatibility issues that can make replicating older analyses on new systems difficult even when the original analysis code is available. Successfully navigating this complexity can often require significant computational know-how, preventing large segments of the biological community from effectively exploring and/or re-using published datasets. We have here addressed this issue in the field of parasitology by implementing paraCell, an interactive cell atlas platform with advanced functionalities and links to external database systems.

paraCell expands the range of experimental results that can be presented by a CELLxGENE atlas to include trajectory inference and tradeSeq, letting scientists easily share more comprehensive versions of their scRNA-seq data. paraCell also integrates CELLxGENE gene sets and incorporates links to a selection of external database systems, streamlining the user experience and enabling cross-system analysis pipelines. Additionally, paraCell adds new analysis and visualization options for both conventional and dual scRNA-seq data, directly increasing the utility of the plugin and addressing the paucity of analysis tools for host-parasite data.

paraCell does all of this within the user-friendly interface provided by CELLxGENE and CELLxGENE VIP, and as such requires no programming ability to use. This lets paraCell serve as a bridge between the biologists who generate single-cell data and the bioinformaticians who often help analyse and annotate it, facilitating the collaborations between wet and dry lab scientists that are now characteristic of single-cell transcriptomics.

paraCell facilitates the generation of novel insights from previously published data, as shown in two of the use-cases presented in this report. In the malaria example, paraCell confirmed existing knowledge and enabled the identification of several novel genes associated with *Plasmodium* gametogenesis, including a protein of unknown function found to be highly upregulated in early development. In the *T. brucei* example, paraCell enabled the detection and analysis of a protein linked to the successful development of the parasite into its replicative form. paraCell can also be used as a platform for the annotation, analysis and visualization of dual scRNA-seq data, further differentiating it from available alternative single-cell browsers (Table 1).

We also present the first attempts to generate and visualise host-parasite. In the *Toxoplasma* example, analysing these novel data and the flexibility conferred by paraCell, we were able to identify ATP6V_0_d2 as significantly upregulated in infected macrophages. ATP6V_0_d2 is a macrophage-specific subunit of the V-ATPase, and upregulation has previously been identified from bulk RNAseq in *Toxoplasma* (e.g. (Li, et al., 2020)); however, it has never been highlighted. In *Leishmania*, the gene is essential for vacuole expansion, likely due to its role in cholesterol biosynthesis (Pessoa, et al., 2019). In the *Theileria* use case, the first Theileria-cow dataset known to date, we detect the differences between an active interferon (IFN) versus a pro-carcinogenic depending on the two hosts. Although those findings are limited to the lack of replicates, they do highlight the possibility of analyzing those datasets. They are unique because of their host-parasite nature and should be helpful for the community in generating hypotheses.

## Conclusion

paraCell is a novel extension of CELLxGENE VIP that incorporates multiple new analysis, visualization, and search options into the plugin, in addition to improving the application’s interoperability with both the underlying CELLxGENE functionality and external database systems. Here we have demonstrated that paraCell enables the generation of new biological insights from published data without requiring any programming ability or highly specialized knowledge on the part of the user, facilitating both the re-use of published datasets in further research and collaboration between bioinformaticians and biologists. Further, the two host-parasite datasets we present here highlight the power of paraCell.

## Methods

### Software Architecture

paraCell makes minimal changes to the server-client architecture established by CELLxGENE VIP, which combines an interactive front-end built via JavaScript libraries such as D3 and jQuery with a Flask backend that implements popular scRNA-seq analysis platforms such as Seurat (v 3.2.3) and Scanpy (v1.6.1) on the server-side (Li, et al., 2022). All this functionality is presented as a plugin built on top of the base CELLxGENE interface.

Instead paraCell utilizes the modular nature of CELLxGENE VIP to extend the plugin, incorporating additional resources on the front-end (e.g. Fuse.js) as well as the back end (e.g. tradeSeq).

paraCell is also available as a containerized application, minimizing set-up requirements for the user.

### Input Data Format

The only input paraCell requires is a properly formatted scRNA-seq AnnData file.

The AnnData file format initially developed by Scanpy is optimized for the storage and manipulation of annotated matrices and has become a popular standard file format within single-cell genomics.

In the context of scRNA-seq, AnnData files combine a central cell-by-gene count matrix (X) with slots holding cell annotations (obsm, obsp), gene annotations (var) and unstructured annotations (uns) into a single data-structure.

CELLxGENE requires input AnnData files that fulfil two criteria - cell barcodes and/or gene names must be unique, and at least one embedding must be available in the obsm slot.

CELLxGENE VIP incorporates an optional text-file as input, letting users display additional information in an atlas (e.g. a description of the dataset) and set initial visualization options (Li, et al., 2022).

paraCell simplifies the user experience and takes full advantage of the annData format by storing all the information necessary to run the plugin in “paraCell_setup” - a basic Python dictionary stored in the uns slot.

This dictionary not only fully replaces the text file used by CELLxGENE VIP, but also controls the activation of core paraCell features. Only features relevant to the dataset in question are activated during set-up, preventing user confusion.

Users are provided with a script capable of inserting the “paraCell_setup” dictionary into a targeted h5ad file - further guidance on the use of “paraCell_setup” is available on the paraCell wiki, here: https://github.com/sii-cell-atlas/paraCell/wiki/Setting-Up-paraCell

### External Database Links

paraCell links cell atlases to external databases: currently available options are NCBI and the VEuPathDB system. The specific database linked to an atlas is established within the “paraCell_setup’’ dictionary.

Multiple paraCell tabs return results in the form of data tables. Any gene names included in such tables are automatically reformatted into hyperlinks, pointing towards the relevant gene profile page on the pre-specified external database system.

### Gene Set Import

Gene sets created in CELLxGENE are automatically made available for import into paraCell, increasing interoperability between the plug-in and the site.

Available gene sets can be selected and imported via a dropdown menu in the *Add Genes* tab.

### paraCell Tabs

The analysis, visualization and utility options provided by paraCell are presented as additional tabs within the original CELLxGENE VIP plugin. paraCell also makes changes to the ‘Add Genes’ tab provided by CELLxGENE VIP, extending its functionality to incorporate links to external database systems and base CELLxGENE gene sets.

### Help

The *Help* tab provides a basic overview of paraCell and its functionality, as well as links to CELLxGENE VIP and the paraCell wiki, which provides comprehensive tutorials on the functionality of each tab.

The paraCell wiki can be found here: https://github.com/sii-cell-atlas/paraCell/wiki

Each individual paraCell tab also includes a link to the corresponding page on the paraCell wiki.

### Download the result files

Beside the graphical interface with different plots, eg in the differential expression analysis, it is possible to download all result as csv files.

### Add Genes

Gene sets created in base CELLxGENE are automatically detected and made available for import into paraCell/CELLxGENE VIP via a drop-down menu in the *Add Genes* tab.

In addition, the *Search Database with User Defined Gene Set* paraCell option made available in *Add Genes* uses available gene sets as search strategies on a pre-specified database system.

### Advanced Gene Search

The *Advanced Gene Search* tab includes two additional search bars, expanding the range of data types that can be used to search a paraCell atlas to include gene names, gene functions and gene ontology terms. Look-up tables for each option are stored within the annData object, which relates the data type to the gene ID indices used by CELLxGENE.

These look-up tables are automatically parsed upon page launch, and the contents of each stored as ordered vectors on the front-end, enabling the quick retrieval of results upon user input. Results are retrieved automatically via a fuzzy search method enabled by Fuse.js.

### Cell Population View

The *Cell Population View* tab utilizes Plotly.js to create interactive scatter graphs of gene expression within a single cell population, split between two conditions. A user-specified gene-level annotation in the AnnData object can then be added to the graph as hover-data.

Results are also available as a jQuery DataTable of differentially expressed genes, obtained by applying the diffxpy (https://diffxpy.readthedocs.io/, version 0.7.4) Welch’s t-test algorithm. The data is filtered to only contain cells belonging to the two conditions being compared prior to the DEG analysis.

### Trajectory Inference

The *Trajectory Inference* tab lets users present pre-computed Slingshot lineages within paraCell.

The coordinates for each lineage are stored as tables in the underlying annData object. The embedding graph used to generate the lineages is indicated via the “pseudoEmbed’’ key of paraCell_setup and is coloured according to the levels of a user-specified categorical annotation.

Novel trajectory inference results can also be generated via the Scanpy implementation of the PAGA algorithm, based on a user specified embedding graph and categorical annotation.

Users are provided with example scripts detailing the preparation of an AnnData object for the *Trajectory Inference* tab in the “slingshot_example” folder provided alongside the main application.

Further guidance on the *Trajectory Inference* tab, as well as a link to the official Slingshot tutorial, is available in the Wiki section of the paraCell GitHub repo.

### tradeSeq

The *TradeSeq* tab enables the presentation of results generated via tradeSeq (v 1.6.0) within paraCell. The tradeSeq R package analyses differential gene expression along trajectories in SingleCellExperiment (Amezquita, et al., 2020) objects.

tradeSeq uses a General Additive Model (GAM) to estimate smooth functions (smoothers) of gene expression along the pseudotime variable of each lineage. tradeSeq is compatible with a wide variety of dimension reduction and trajectory inference methods and is applicable to both simple and complex trajectories (Van den Berge, et al., 2020). Rpy2 (v3.3.5) allows R scripts to run embedded in a Python process, enabling the paraCell backend to interact with tradeSeq.

tradeSeq includes several tests capable of identifying differential gene expression patterns both within and between lineages. paraCell can present pre-computed results generated via associationTest(), a tradeSeq package function which assesses whether gene expression is associated with pseudotime either globally, or for each specific lineage.

Test results are extracted from the SingleCellExperiment object tradeSeq operates on, stored as a table in the AnnData object, and then presented as a jQuery DataTable within the *TradeSeq* tab.

paraCell can also plot the smoothers estimated by tradeSeq, visualizing the relationship between a given gene and the progression of specified lineages. This function requires that results of the GAM fitting process are extracted from the SCE data object, split into columns, and saved into the annData object as a collection of vectors.

paraCell provides users with scripts that extracts the necessary objects from a given SingleCellExperiment object containing tradeSeq results and adds them to a specified AnnData file.

paraCell provides users with scripts that automate both the extraction of tradeSeq results from a given SCE data object, and the addition of those results to a user specified AnnData object, streamlining the use of the *tradeSeq* tab.

Further guidance on the *tradeSeq* tab, as well as a link to the official *tradeSeq* tutorial, is available in the Wiki section of the paraCell GitHub repo.

### Host-Parasite Interactions

The *Host Parasite Interactions* tab is designed for the analysis of dual scRNA-seq datasets in which each cell contains both host and parasite genes.

The Host-Parasite Violins option generates a multi-panel violin plot indicating the percent concentration of host or parasite genes within each level of a specified categorical annotation.

This requires that the percent host and parasite content for each cell is pre-calculated and stored as a cell level annotation in the annData object.

Host and parasite UMAPs are generated by subsetting the annData object to contain exclusively host or parasite genes, before reducing dimensions and clustering via Scanpy.

Host/parasite subsetting relies on premade vectors listing every host and every parasite gene in the dataset respectively (host_genes, parasite_genes), stored in the AnnData object.

Both UMAPs are presented as interactive scatter graphs via Plotly.js. Cell selections made via the Plotly.js lasso select function on one graph (e.g. Host UMAP) automatically applies one of two update options to the opposing graph (e.g. Parasite UMAP).

The “Highlight Cells” update option alters the traces on the opposing graph, colouring cells with the same cell IDs as the selected cells red and all other cells blue.

The “Recluster” update option regenerates the opposing graph, this time subsetting the data object to only contain the selected cells in addition to the original host/parasite gene subset.

Host/Parasite specific gene markers are generated for the levels of specified categorical annotation by subsetting the annData object to contain only host or parasite genes, before DEG analysis via the t-test, t-test with overestimated variation, or Wilcoxon Rank Sum options provided by Scanpy.

An example use-case demonstrating the generation and preparation of a host-parasite AnnData object for paraCell can be found in the “host_parasite_example” folder provided alongside the main application.

## Data Generation

### *Toxoplasma gondii* Tachyzoite Cell Culture

Wildtype *T. gondii* tachyzoites were grown at 37°C with 5% CO_2_ in human foreskin fibroblasts (HFFS) cultured in Dulbecco’s Modified Eagle’s medium (DMEM) (ThermoFisher Scientific) supplemented with 3% FBS, 2mM L-glutamine and penicillin-streptomycin.

### Mice

C57BL/6 (B6) mice (JAX 000664) purchased from Jackson Laboratories. All mice were female, aged between 6-12 weeks. Procedures were conducted in accredited animal facilities under the project license PPL30/3423, authorized by the UK Home Office Animal Procedures Committee.

### BMDM Isolation and Infection with *Toxoplasma gondii*

6–12-week-old B6 mice were euthanized and femur and tibia bones were pooled from 2-3 mice. Tissue was removed from the bones using forceps, bones were placed in ethanol for 30 seconds to sterilise and then transferred to 1X sterile PBS. Bone marrow was extracted and resuspended in 1 mL ACK lysis buffer (ThermoFisher Scientific). Cells were then incubated in ACK lysis buffer for 2-5 minutes before washing. BMDMs were cultured in RPMI (ThermoFisher Scientific) complete media, supplemented with 10% filtered FBS, 2 mM of L-glutamine, penicillin-streptomycin and 50 µM ß-mercaptoethanol (ThermoFisher Scientific). Cell were diluted to 5×10^5^ cells/mL and 30ng/mL of M-CSF (Biolegend) was added to promote BMDM differentiation. Culture media was replaced with fresh complete RPMI and 30ng/mL M-CSF 3-4 days following isolation.

After 6 days of differentiation, BMDMs were infected with wildtype RHΔKu80 *T. gondii* tachyzoites at an MOI of 1.5, in the presence or absence of 20 ng/mL murine recombinant interferon-γ (IFN-γ) (Biolegend). Cells were incubated at 37°C with 5% CO_2_ for 24 h before harvesting for scRNA sequencing.

### BMDM Sample Preparation for 10X Genomics scRNA Sequencing

Briefly, cells were washed twice with sterile PBS, then incubated with 4 mL of trypsin at 37°C with 5% CO_2_ for 5-7 minutes. 6 mL of complete RPMI media was then added to gently resuspended the cells. Cells were spun down at 1500 x g for 5 minutes, resuspended in 10 mL complete RPMI media and then filtered using a 40 µM filter before diluting in 0.2% Trypan Blue and counting. For 10X scRNA sequencing analysis, 1.6 x 10^4^ cells/condition were resuspended in 0.04% BSA in PBS. Cells were counted again and loaded onto the 10X Genomics technology for single cell RNA-sequencing.

### Single cell preprocessing

The raw fastq files were mapped with Cellranger (version 7.0.0) against a combined reference of *Toxoplasma gongii* (version 59 VEuPathDB) and Mouse (version GRCH38_mm10_ensemble93). With custom scripts, we extended the parasite’s UTR by 2.5kb. The read count matrix was analysed with Seurat (version 3) following the integration tutorial. As parameters we chose: Cells must have between 200 – 5000 expressed genes, default Seurat integration, 30 PCA dimension and a resolution of 0.5. The resulting object was saved as AnnObject for paraCell.

### Cow-Theileria

Two *Theileria* infected cell lines were utilised. One line (Sah1 (82H)) represented an *ex vivo* isolate derived from an infected calve of the Sahiwal breed (*Bos indicus)* while the other (Hol 3 (12886)) was derived from an infected Holstein calf (*Bos taurus*), as reported (Larcombe, et al., 2022) . Both Sahiwal and Holstein animals were infected with sporozoites from the same, *T*. *annulata* Hissar stock and cultured as standard in RPMI-1640 medium (Sigma-Aldrich) supplemented with 10% FCS, 4 mM L-glutamine and 50 μM β-mercaptoethanol, at 37 ^0^C for a limited period (l2 passages). Both cell lines were fully established based on the level of infection (>95% macroschizont infected cells).

Cultures were set up at 1X10^5^cells/ml and cultured for 48 hours. Cell counts and viability were determined by trypan blue exclusion using a Countess^TM^ 3FL automated cell counter (Invitrogen). Cell viability was 94% (Sah 1) and 95% (Hol 3), while growth for the Holstein line was 1.58 fold greater than the Sahiwhal.

Multiome nuclei extraction protocol was adapted from(dx.doi.org/10.17504/protocols.io.rm7vzyx75lx1/v1), cells were harvested by centrifugation (400 g for 10 min), and lysed for 10min on the rotor at 4 C with 1X CST lysis buffer (as indicated in dx.doi.org/10.17504/protocols.io.rm7vzyx75lx1/v1). Lysate was filtered with 40 μm mini cell strainer into a clean pre-labelled 5.0 mL Eppendorf tube and PBS + 1% BSA was added to dilute the lysis buffer. The tube was spun down 500g, 4°C, 5 min. Nuclei were resuspended and counted with Neubauer haemocytometer; 10,000 nuclei were loaded for each chip line on a 10X Genomics Multiome J chip (#1000230). Libraries were prepared according to the 10X Genomics multiome protocol (#1000285, #1000215, #1000212) and sequenced at Glasgow University Polyomics facility on an Illumina NextSeq 2000.

### Single cell preprocessing

The raw fastq files were mapped with Cellranger (version 7.0.0) against a combined reference of *Tannulata Ankara* (version 59 VEuPathDB) and *Bos taurus* genome from Ensembl (version CowARS-UCD1.2_UTR). With custom scripts we extended the UTR of the parasite by 2.5kb. The read count matrix was analysed with Seurat (version 3) following the integration tutorial. As parameters we chose: Host cells must have between 1000 – 75000 expressed genes, and maximal 5000 parasite UMI, default Seurat integration, 30 PCA dimension and a resolution of 0.5. The resulting object was saved as AnnObject for paraCell. Differentially expressed genes were found with Welch’s test (cellxgene), FDR < 0.05, absolute log2(FC) > 0.6; GSEA was performed using database C2.all.v7.2 with default parameters.

## Declarations

### Availability of Data and Materials

All the datasets used to demonstrate paraCell functionality can be found as publicly available cell atlases, as listed in the following table. **For best performance, the Chrome browser should be used.**

| Organism | URL | Reference |
| --- | --- | --- |
| *Plasmodium berghei* | https://cellatlas-cxg.mvls.gla.ac.uk/Plasmodium_berghei-Multi.Tissue/ | Hentzschel et al (Hentzschel, et al., 2022) |
| *Trypanosoma brucei* | https://cellatlas-cxg.mvls.gla.ac.uk/Trypanosoma_brucei-slender2stumpy/ | Briggs et al (Briggs, et al., 2021) |
| *Toxoplasma-Mouse* | https://cellatlas-cxg.mvls.gla.ac.uk/Toxoplasma_gondii-murine.bone.marrow.derived.macrophages/ | Novel dataset |
| *Theileria-Cow* | https://cellatlas-cxg.mvls.gla.ac.uk/Theileira_annulata-Bos_indicus_taurus_macrophage/ | Novel dataset |

The newly generated raw fastq data are being submitted to ENA. Upon acceptance of the paper, the AnnObject will be publicly accessible.

### Code Availability

paraCell is an open-source project and MIT licensed application, available here: https://github.com/sii-cell-atlas/paraCell/

A Docker Image of paraCell is also available at: https://hub.docker.com/r/edwardagboraw/paracell

### Tutorial

Comprehensive guidance on the installation and usage of paraCell is available on the paraCell wiki, here: https://github.com/sii-cell-atlas/paraCell/wiki

### Authors’ contribution

EA implemented paraCell, performed bioinformatics analysis and wrote the first draft. WHH tested the implementation and maintains the paraCell server. FH performed the malaria analysis and wrote that section. EB performed the *Trypanosoma* analysis and wrote that section. DS and BS performed the *Theileira* experiment, analysis and wrote that section. AH and DA perform the *Toxoplasma* experiments. CRH performed the *Toxoplasma* analysis and wrote that section. KC assisted with connecting paraCell to VEuPathDB and gave general feedback on implementation. TDO conceptualised the project, organised funding and wrote the paper. All authors gave feedback on the manuscript and agreed with it.

## Acknowledgements

We would like to thank Scott Arkison (School of Infection and Immunity) for maintaining the computational infrastructure that hosts paraCell.

