## Supplemental material for "paraCell: A novel software tool for the interactive analysis and visualization of standard and dual host-parasite single cell RNA-Seq data"

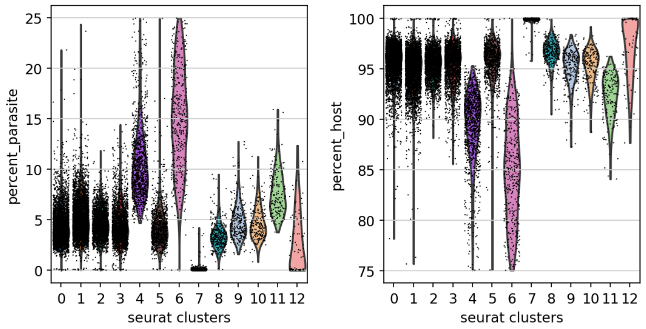


Supplemental Figure 1: Violin plots showing the frequency of mRNA in Cow versus *Theileria* for the different clusters.

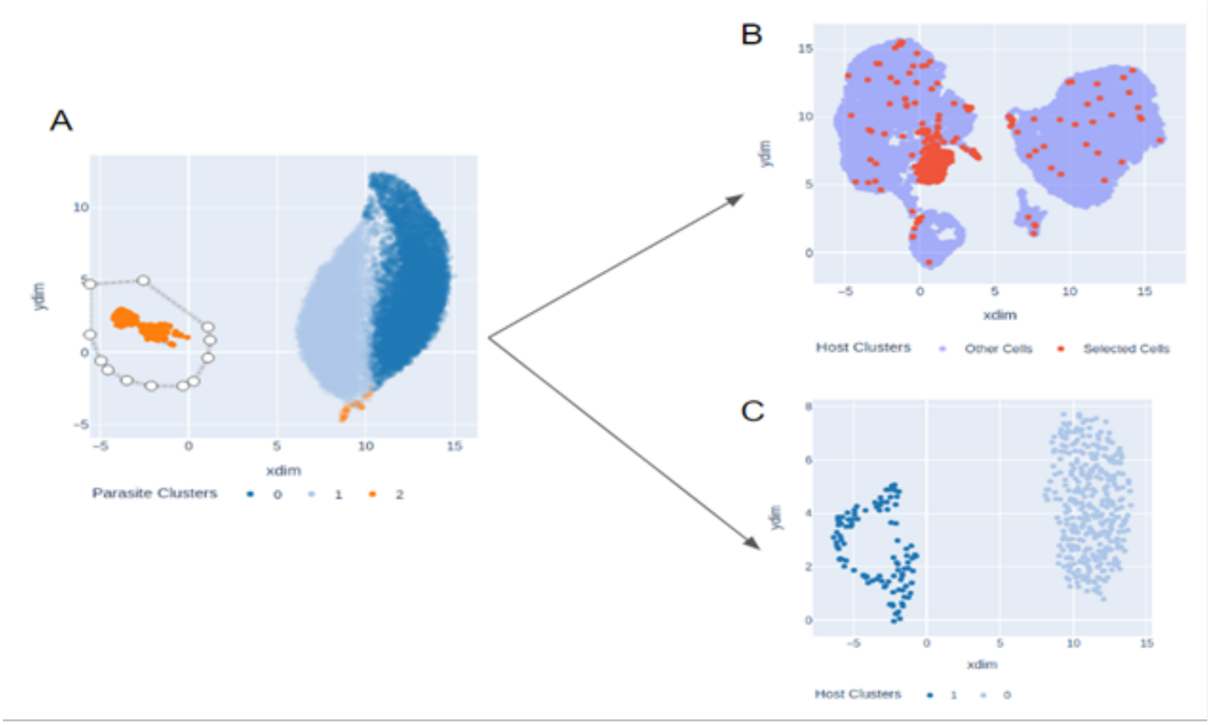


Supplemental Figure 2: [A] Parasite UMAP presented within the Host-Parasite Interaction tab. A cluster of cells has been selected via the Plotly lasso tool. [B] The Host UMAP is updated so that the cells selected on the Parasite UMAP are highlighted in red and all other cells are blue. [C] Host UMAP produced by subsetting the data object to only contain host genes, subsetting again to only include the cells selected on the parasite UMAP (as indicated in A), and then clustering the data and generating the UMAP.
